# Asymmetric Spring and Autumn Phenology Control Growing Season Length in Temperate Deciduous Forests

**DOI:** 10.1101/2025.10.21.683688

**Authors:** Xiaojie Gao, Mark A. Friedl, Andrew D. Richardson, Valerie J. Pasquarella, John O’Keefe, Amey S. Bailey, Jonathan R. Thompson

**Affiliations:** Harvard Forest, Harvard University, MA, USA; Department of Earth and Environment, Boston University, MA, USA; Center for Ecosystem Science and Society, Northern Arizona University, AZ, USA; School of Informatics, Computing, and Cyber Systems, Northern Arizona University, AZ, USA; USDA Forest Service, Hubbard Brook Experimental Forest, NH, USA

## Abstract

Forest phenological responses to climatic variation among species and populations across broad spatial scales remain poorly understood. Here, we quantify four decades of phenological dynamics for 10 major deciduous tree species by compiling a unique dataset that integrates high spatial resolution remote sensing with extensive field inventory data across the Northeastern and Midwestern United States. We found that spring and autumn phenology impose asymmetric controls on growing season length, with spring regulating interannual variation and autumn driving long-term trends. Spring phenology across all species was primarily controlled by temperature, while autumn phenology was influenced by spring phenology and an interacting suite of species-specific environmental factors. Our study demonstrates the promise of combining high resolution remote sensing with forest inventory data to investigate phenological dynamics over large regions. More importantly, our results provide new insights into how species-specific sensitivities to environmental drivers regulate growing season length in temperate forests.

## Introduction

Leaf phenology regulates the seasonality and duration of forest photosynthesis and controls biosphere-atmosphere exchanges of carbon, water, and energy ^1–4^. A large body of evidence demonstrates that climate change is altering forest phenology at regional-to-global scales ^2,5,6^. These changes have far-reaching consequences for community composition and structure ^7,8^, phenological synchrony across trophic levels ^8–10^, and biosphere-atmosphere interactions that create both positive and negative feedbacks to the climate system ^2,11–13^. Because forests provide the largest sink for carbon in the terrestrial biosphere ^14,15^, understanding the response of forest phenology to environmental drivers is essential for understanding how forests interact with the climate system and for predicting how forest ecosystems are likely to change in the coming decades.

While there is broad consensus that climate change is altering forest phenology, recent studies have demonstrated that species-level responses of phenology to environmental drivers are heterogeneous and poorly understood ^16–18^. Numerous studies have examined the phenological response of individual species or small cohorts of species to climate change ^19–21^. Similarly, broad-scale patterns at the level of plant functional types are widely documented ^2,22^. However, comprehensive understanding of how species-specific responses to climate variability aggregate to control phenology at the scale of multiple ecoregions is lacking.

This knowledge gap persists partly due to data limitations. The Pan European Phenological (PEP725) database ^23^ provides species-level information and has been extensively used for investigating phenological shifts ^21,24,25^. Unfortunately, the PEP725 data are limited in regard to climate, geographic area (only in Europe), and the range of species that have long-term phenological records. In the United States, the National Phenology Network (US-NPN) dataset ^26^ includes a wide range of species and climate types, but the US-NPN time series is too short to support assessment of trends and drivers of change. Both PEP725 and the US-NPN suffer from variable data quality because the observations are collected by volunteers rather than experts. Satellite remote sensing has been widely used to capture phenological dynamics at continental-to-global scales from multiple sensors, some of which now span multiple decades ^27–29^. However, currently available datasets derived from satellite remote sensing are either too coarse in spatial resolution (500-8,000 m; e.g. ^30,31^) or too short (<10 years; e.g. ^32,33^) to be used to analyze species-specific phenological dynamics. Because phenology reflects an organismal response to environmental forcing but the impact of phenological changes occurs at the scale of ecosystems, it is important to improve understanding of how species-level responses to climate variability and change contribute to ecosystem-scale dynamics in phenology and ecosystem function.

To address this need, here we use a novel dataset that spans four decades (1985-2023) and includes 108,264 plot years of species-specific phenological records for 10 major tree species across the Northeastern and Midwestern United States, a region which includes the dominant species of the vast eastern temperate deciduous forest of North America (**Extended Data Figure 1**; **Extended Data Table 1**). This dataset combines ground measurements from 2,776 Forest Inventory and Analysis (FIA) plots with satellite-observed phenology at high spatial resolution (30 m). Supplemented with long-term (> 30 years) in-situ measurements of phenology collected at two research forests (**Extended Data Figure 1**), we quantify the nature and magnitude of species-level changes in spring and autumn phenology, as well as their contributions to dynamics in growing season length. Our analysis focuses on two core questions: how does climate forcing control growing season length, and how is climate change driving species-specific phenological responses across temperate forests of the Northeastern and Midwestern United States? Since different components of phenological variation cause distinct ecological responses ^34^, we investigate three dimensions of phenological change across time in our data set: (1) long-term trends in the timing of spring and autumn phenology; (2) interannual variation in the timing of phenology across the study period; and (3) stationarity in interannual variation in spring and autumn phenology across the study period. Using historical climate data, we estimate Bayesian hierarchical models to quantify how species-specific sensitivity of phenology to environmental drivers explains observed patterns in the growing season length of temperate forests.

## Results

### Asymmetric contributions of seasonal phenology to GSL dynamics

Over the past four decades, ∼5-35% of plots across our study region showed statistically significant advancing or delaying trends (*p* < 0.05) in the remotely sensed start-of-season (SOS), end-of-season (EOS), or growing-season-length (GSL) for each tree species included in our analysis (**Fig. 1a; Extended Data Figure 3**). Across sites with significant trends, linear models and methods that are robust to outliers and account for heteroskedasticity both estimate median trends of ∼0.2-0.3 days yr^-1^ across all species for the three metrics (SOS, EOS, and GSL) (**Extended Data Figure 3**), equivalent to mean changes of 8-12 days across the ∼40-year study period.

**Figure 1.**
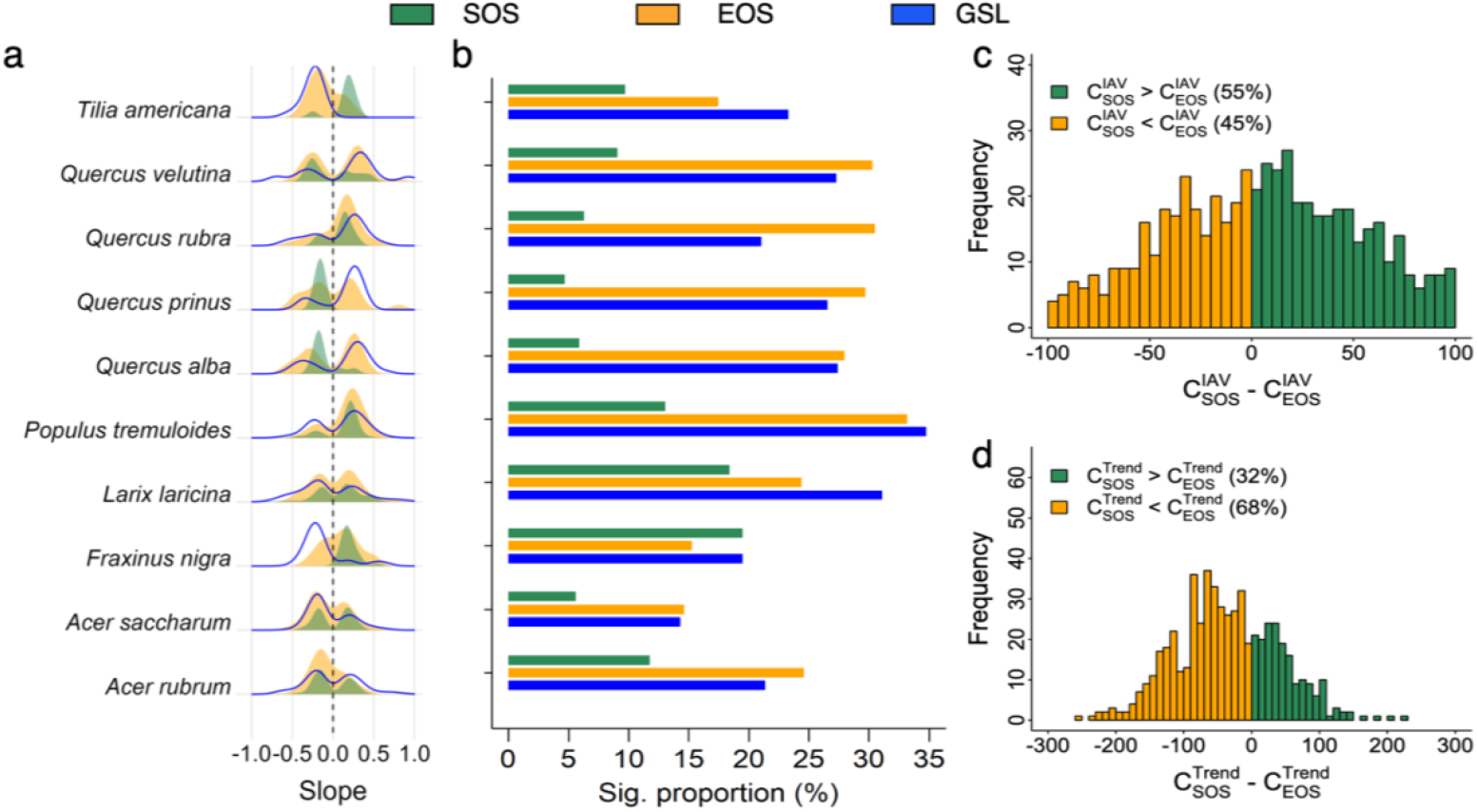
Phenological dynamics derived from remotely sensed phenology for each species. **a)** Distribution of significant trends (*p*<0.05) in start-of-season (SOS), end-of-season (EOS), and growing season length (GSL) for each species across all sites. **b)** The proportion of sites with statistically significant trends for each metric and species. **c)** and **d)** show the frequency distribution for sites where SOS explains a larger proportion of interannual variation (IAV) in GSL relative to EOS 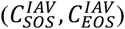, and where SOS and EOS explain a larger proportion of long-term trends in GSL 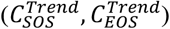. The percentage values in parentheses show the percentage of sites for each case. Note that although 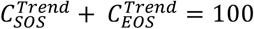, their values can exceed 100 or be less than 0, and thus their difference is not bounded to [-100, 100] (See Methods).

Trends in GSL are more strongly controlled by trends in EOS than by trends in SOS. Qualitatively, this is evident in **Figure 1a** and **Extended Data Figure 3**, which show that (with exception of *Fraxinus nigra*) the distribution of statistically significant GSL slopes is more closely aligned with the distribution of EOS slopes than with the distribution of SOS slopes. The dominant role of EOS in regulating GSL trends is also supported by the proportion of statistically significant trends in each phenometric (**Fig. 1b; Extended Data Figure 3**); with the exception of *F. nigra*, the proportion of statistically significant EOS trends across all sites (∼15%-35%; **Fig 1b**) aligns closely with the proportion of statistically significant GSL trends. And, across all species, the proportion of statistically significant trends in both metrics was consistently higher than the proportion for SOS trends (∼ 5%-20%).

To explore this further and quantify the relative contribution of variability in SOS versus EOS to interannual and long-term dynamics in GSL, we decomposed the time series for each phenological metric into two components: the first quantifies the magnitude of interannual variation over the study period and the second quantifies long-term trend (See Methods). Our results show that, for sites exhibiting statistically significant trends in GSL, SOS was the dominant control on interannual variation in GSL at 55% of the sites, whereas EOS was the dominant control on long-term trends in GSL at 68% of the sites (**Fig. 1c**). This pattern was generally consistent across all sites and species, especially the stronger role of EOS in regulating GSL trends (**Extended Data Figure 4**). These results identify the unique contributions of SOS versus EOS to interannual variation and long-term trends in GSL: interannual variation in SOS dominants interannual variation in GSL, while long-term trends in GSL are controlled by shifts in EOS.

To evaluate the robustness of the results described above, which are derived from remote sensing, we applied the same methodology to long-term in-situ phenology records collected for individual trees at two research forests located in the Northeastern United States (**Extended Data Figure 1**). We found patterns in the in-situ measurements that further support our results and conclusions derived from remote sensing (**Extended Data Figure 5; Extended Data Figure 6**). Specifically, relatively few trees showed statistically significant trends in the in-situ measurements of SOS. In contrast, numerous species showed statistically significant trends in both EOS and GSL. The in-situ measurements also exhibited interannual variation and trends in SOS, EOS, and GSL that mirror the patterns we identified in the remote sensing-based results.

### Changes in Interannual Variation of Phenology

To quantify the stationarity of interannual variation in each phenometric, we detrended the times series of SOS, EOS, and GSL and used a moving window approach to evaluate whether interannual variation in each metric was stable across the study period (**Fig. 2a**; See Methods). Our results reveal widespread non-stationarity in the interannual variation of phenology across our study region. 51% of the sites exhibited statistically significant trends (*p* < 0.05) in the interannual variation of SOS, with 24% exhibiting increases in interannual variation and 27% exhibiting decreases over the past four decades (**Fig. 2a**). In contrast, 54% of sites exhibited increases in interannual variation for EOS, while only 11% showed significant decreases (**Fig. 2b**), suggesting a strong and widespread increase in the magnitude of interannual variation in autumn phenology. Similar to EOS, 40% of sites showed significant increases in the interannual variation of GSL and 18% showed significant decreases (**Fig. 2c**). This lack of stationarity in interannual variation is widespread across sites and species and is present in both the remotely sensed and in-situ phenology measurements (**Extended Data Figure 7; Extended Data Figure 8**).

**Figure 2.**
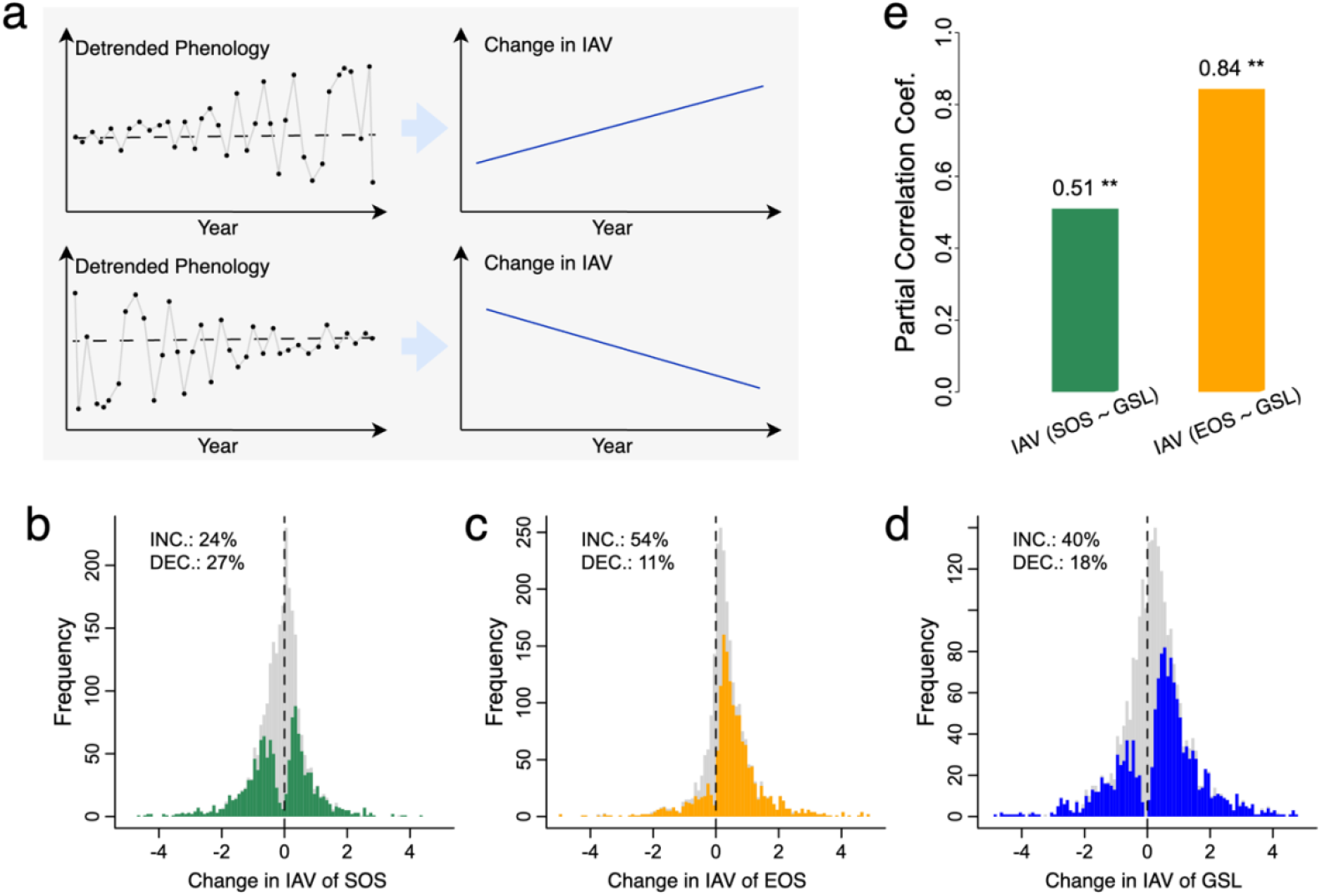
Stationarity, or changes in interannual variation (IAV), of phenology. **a)** schematic showing detrended phenological time series and corresponding hypothetical trends in IAV. **b), c)**, and **d)** show the distribution of trends in IAV across sites for SOS, EOS, and GSL, respectively. The percentage of statistically significant (*p*<0.05) increases (INC) and decreases (DEC) are included for each plot. **e)** partial correlation coefficients for IAV trends between SOS and GSL and between EOS and GSL, respectively, across sites. The number on top of each bar shows the partial correlation coefficient value and ** indicates that the partial correlation is significant (*p*<0.05).

Changes in the interannual variation of SOS and EOS across the study period can either amplify or diminish stationarity in GSL, depending on whether their changes are correlated. To quantify the relative contribution of SOS and EOS to changes in interannual variation of GSL, we estimated the partial correlation between the trend in interannual variation for GSL and the corresponding trends in SOS and EOS, respectively, whiling controlling for the other. We found that the partial correlation between the trend in interannual variation of EOS and the trend in interannual variation of GSL was ∼1.6x higher than the corresponding partial correlation between GSL and SOS (0.84 *vs*. 0.51; **Fig. 2e**), suggesting changes in interannual variation of EOS had a larger effect than corresponding changes in SOS on the stationarity of interannual variation in GSL.

### Drivers of Phenological Variation

To investigate the bioclimatic drivers of SOS and EOS at both the site and species level, we transformed the values for each phenometric into site-specific anomalies (z-scores). We then used Bayesian hierarchical models to predict site-specific anomalies in SOS and EOS as a function of seasonal variation in temperature, precipitation, and solar radiation. The resulting models quantify the sensitivity of phenology to interannual variation in these forcing variables, controlling for site-level characteristics and species. We selected the best models in terms of structure and accuracy and standardized the predictors into site-specific z-scores, which allowed us to evaluate the relative contribution of each predictor to SOS and EOS (See Methods).

Unsurprisingly, SOS across all species was most strongly controlled by spring temperature (R^2^ = 0.37, *p* < 0.05; **Fig. 3a; Extended Data Figure 9**). Compared to temperature, the relative controls of springtime precipitation and solar radiation were negligible. Six out of ten species (*Quercus velutina, Quercus prinus*, and *Quercus alba, Larix laricina, Acer saccharum*, and *Tilia americana*) showed statistically significant responses to precipitation and nine species (except for *Larix laricina*) showed statistically significant responses to solar radiation, but the magnitude in each case was far lower than the effect of temperature. Indeed, the heterogeneous and weaker trends in SOS were consistent with dynamics in spring temperature in this region over the study period, which also showed heterogeneous and weak trends (**Extended Data Figure 10**).

**Figure 3.**
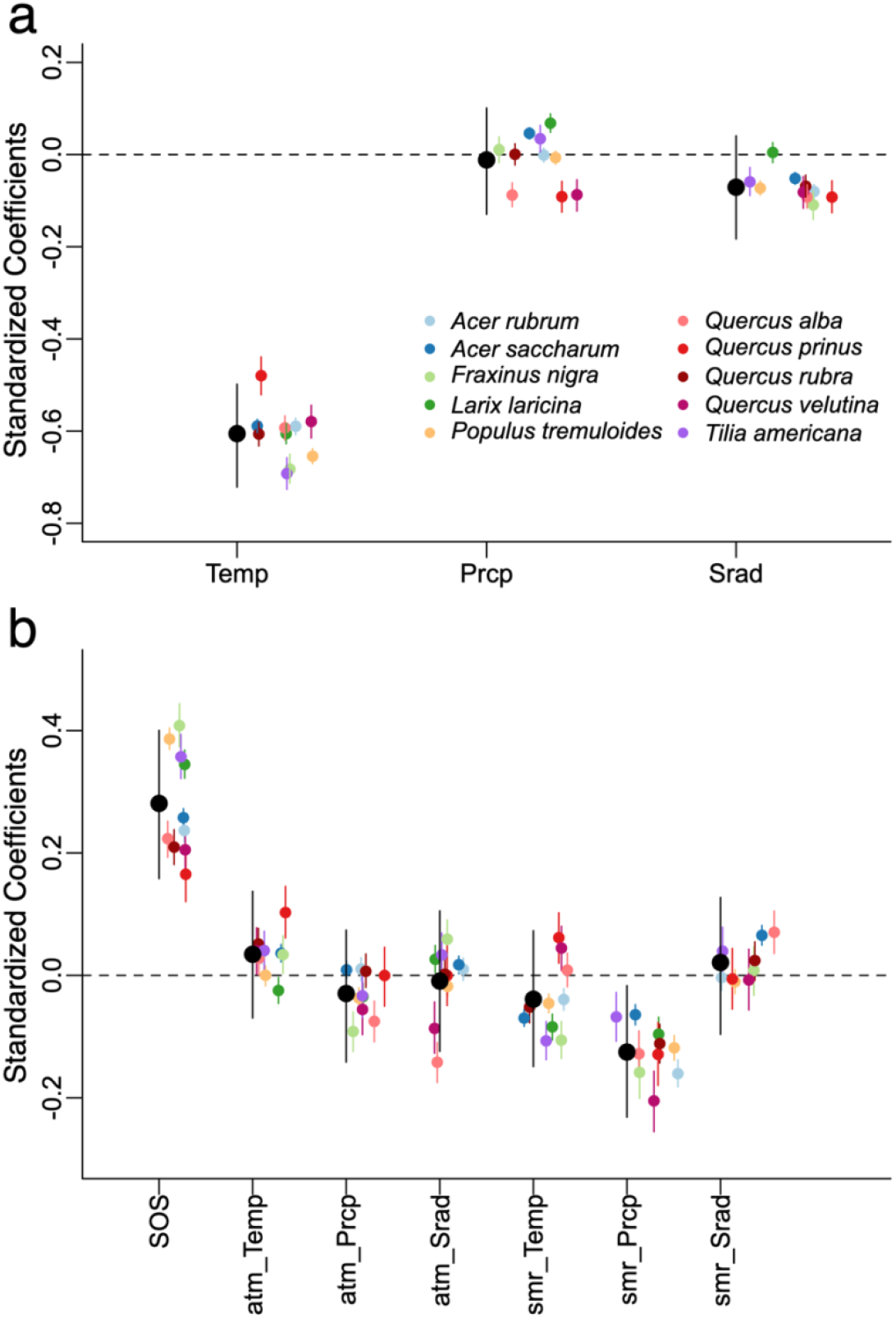
Standardized coefficients for spring (a) and autumn (b) phenology models. Black points show the average effect, and colored points show species-level effects. Bars represent 95% credible intervals. Seasonal climate factors include temperature (Temp), precipitation (Prcp), and solar radiation (Srad). For spring phenology, the seasonal climate factors were calculated as their mean values during March–May. For autumn phenology, mean climate factors were calculated for summer (smr) during June–August and for autumn (atm) during September–November. The effect of spring phenology (start-of-season, SOS) on autumn phenology was also included.

Compared to SOS, the drivers of EOS were more heterogeneous across species (R^2^ = 0.19, *p* < 0.05; **Fig. 3d**; **Extended Data Figure 9**). Controlling for other predictors, SOS was positively correlated with EOS, i.e., earlier (later) SOS was associated with earlier (later) EOS. Greater summer precipitation was correlated with earlier EOS across all species, a result that has also been reported in recent studies with a small set of species ^35,36^. Other predictors, including autumn temperature, autumn precipitation, autumn solar radiation, summer temperature, and summer solar radiation, exerted species-specific control on EOS. For example, higher autumn precipitation was significant for five of the ten species. With the exception of *Quercus velutina* and *Quercus prinus*, warmer summer temperatures were also associated with earlier EOS, while warmer autumn temperatures were generally associated with later EOS. Summer and autumn temperatures had opposing effects on EOS, which is consistent with results from recent studies ^37,38^. Overall, these results show that the drivers and mechanisms that control the timing of EOS differ across species.

## Discussion

Results from previous studies that rely on in-situ observations to investigate long-term phenological dynamics provide insights into species-specific responses to climate change, but are constrained by geographic limitations inherent to available phenological records ^21,39,40^. Remote sensing-based studies, in contrast, cover large areas but lack the spatial resolution needed to reveal species-specific responses ^2,27,41^. Our study overcomes these constraints by combining forest inventory measurements with high spatial resolution remote sensing to create a novel phenology dataset that is simultaneously long-term, geographically extensive, and species-specific. This framework provides a unique foundation for exploring species-specific trends and controls on long-term phenology in temperate forests and is readily transferable to anywhere forest survey data exist. Drawing on four decades of phenological records for 10 dominant deciduous tree species across the Northeastern and Midwestern United States, along with long-term in-situ data collected by experts at two research forests, our analysis reveals important and previously undocumented phenological dynamics and sensitivities to environmental change.

Specifically, results from our analysis demonstrate that both autumn phenology and GSL exhibit statistically significant trends over the last four decades, while long term trends in spring phenology are relatively uncommon across our study region. The absence of significant trends in spring phenology that we document contradicts results from previous studies based on shorter records or coarser resolution remote sensing concluding that the timing of spring phenology is advancing in temperate forests of North America ^2,22,42,43^. However, the lack of advancement in remotely sensed spring phenology that we identify is consistent with the absence of trends in long-term in-situ phenology records (**Extended Data Figure 5–6**) and with long-term dynamics in the main driver that controls spring phenology (temperature, **Fig. 3; Extended Data Figure 10**). In contrast, systematic warming during summer and autumn (**Extended Data Figure 10**) did not yield homogeneous shifts in autumn phenology across species and sites, even though summer and autumn temperatures were shown to exert statistically significant controls on autumn phenology for the majority of species included in our analysis (**Fig. 1**; **Fig. 3**; **Extended Data Figure 3–6**). Instead, trends in autumn phenology were controlled by the combined effects of multiple climate forcings (**Fig. 3**; **Extended Data Figure 11**). Indeed, our results identify heterogeneous species-specific responses in autumn phenology to individual environmental factors (**Fig. 3**) ^44,45^. Hence, the same environmental conditions can yield distinct responses in the timing of autumn phenology across species.

Moreover, we show that the timing of leaf emergence in spring is the dominant factor regulating interannual variation in GSL, but the timing of senescence controls long-term trends and whether or not interannual variation in GSL is stationary (**Fig. 1; Fig. 2**). These asymmetric dynamics not only have important implications for seasonal and long-term variations in ecosystem function ^3,9,10,46,47^, but also reflect the impact of climate change on forest ecosystems. For instance, accurate representation of interannual variation in GSL is crucial for capturing interannual dynamics in ecosystem carbon uptake ^12,48^. Our results show that a large proportion of interannual variation of GSL is explained by interannual variation in spring phenology, whereas systematic shifts in the timing of autumn phenology over multiple decades are the key driver of systematic changes in GSL over the long term (**Fig. 1**). Therefore, models that capture short-term interannual variation in GSL but ignore autumn phenology dynamics are unlikely to provide reliable forecasts of ecosystem carbon uptake under future climate scenarios. More generally, our results demonstrate that interannual variation and long-term trends in GSL are controlled by asymmetric and season-specific phenological shifts that respond to species-specific environmental drivers, underscoring the distinct roles of spring and autumn phenology in controlling GSL dynamics.

Our results identify a strong correlation between interannual variation in spring and autumn phenology across all species (**Fig. 3**), which explains why interannual variation in GSL is controlled by spring phenology rather than autumn phenology (**Fig. 1**). Spring and autumn phenology are generally modeled independently of each other, even though experimental ^49,50^ and observational studies ^40^ have noted their correlation. Although the mechanisms driving the spring-autumn phenological correlation remain debated ^40,49–53^, our results indicate that spring phenology affects interannual variation in GSL both directly and indirectly, the latter through shifts in autumn phenology. This result implies that autumn phenology may serve as an integrative indicator of response of deciduous forests to climate change. Hence, interannual variation in GSL reflects variability in spring phenology—and thus spring temperature effects—while long-term trends and stationarity in GSL reflect the impact of climatic change-induced changes in multiple environmental drivers late in the growing season.

Together, our results advance current understanding in three aspects (**Fig. 4**). First, contrary to the widespread expectation that spring phenology has significantly shifted in the Northeastern and Midwestern United States, we show that autumn phenology and GSL, rather than spring phenology, exhibit more pronounced trends over the last four decades. Second, although GSL measures the time period between spring and autumn phenological events, these variation in spring and autumn phenology do not contribute equally to dynamics in GSL. Instead, we find that spring and autumn phenology exert asymmetric controls on GSL, with spring regulating interannual variation and autumn driving long-term trends and stationarity. Third, while spring phenology primarily responds to temperature, autumn phenology is shaped by an interacting suite of environmental factors that vary by species and is strongly influenced by the timing of spring phenology across species.

**Figure 4.**
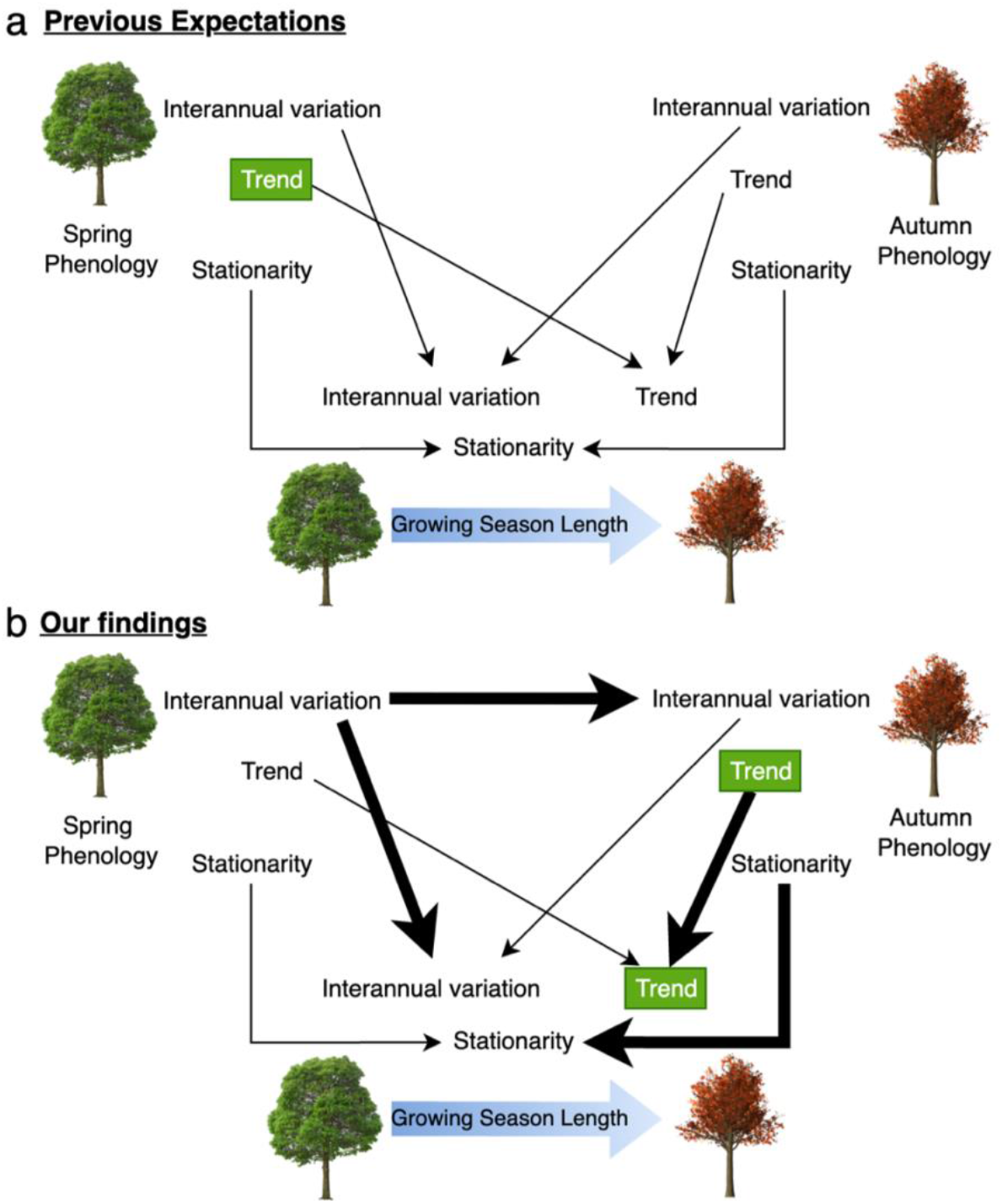
Conceptual model comparing previous expectations (a) and our findings (b) on the seasonal phenological controls of growing season length dynamics. Black arrows represent the direction and relative strength of effects, with arrow width indicating effect magnitude. Green “Trend” labels denote statistically significant long-term phenological trends. In contrast to current understanding (a) that spring and autumn phenology are independent and contribute equally to the dynamics in growing season length, our findings (b) suggest strong intra-annual coupling between spring and autumn phenology and asymmetric contributions to interannual variation, long-term trends, and stationarity in growing season length.

In conclusion, our analysis demonstrates that combining long-term phenological observations from high spatial resolution satellite imagery with ground observed species composition opens new opportunities to investigate long-term and species-specific phenological dynamics and responses to climate change over large geographic regions. More importantly, our findings highlight species-specific sensitivities in seasonal phenological responses to environmental drivers that regulate the length of the growing season in temperate deciduous forests.

## Methods

### Study sites with dominant species

We used 22,658 Forest Inventory and Analysis (FIA) sites with stable dominant deciduous tree species surveyed during 1998-2023 across 20 states in the Northeastern and Midwestern U.S. (Region 9 in the level 1 of the Ecological Regions of North America defined by the U.S. Forest Service) ^54^. The stable dominant species, if present, for an FIA site was determined by three criteria: 1) the basal area of the species must exceed 60% of the site area; 2) the site must be measured more than twice with at least a 10-year interval and the recent measurement must be within the last 10 years, i.e., 2013-2023; and 3) among the measurements in different years, the dominant species must experience no change. The precise FIA plot locations were obtained through a memorandum of understanding between the U.S. Forest Service and Harvard University (Data Use Agreement 19-MU-11242305-016). We aggregated species-specific basal area in FIA subplots to the site level by taking the average and used the central subplot location to represent the site location. We removed FIA sites with non-forest subplots, with no stable dominant species, and with dominant species that occupied fewer than 50 sites (this threshold was determined by analyzing the distribution of dominant species and their number of sites), resulting in 10 major species across 2,776 FIA sites (**Extended Data Figure 1; Extended Data Table 1**).

### Four decades of forest phenology from Landsat

We obtained surface reflectance time series of Landsat 4, 5, 7, 8, 9 Analysis Ready Data (Collection 2) ^55^, and the Harmonized Landsat Sentinel-2 (HLS) ^56^ at 30 m spatial resolution during 1984-2023 for all the selected FIA sites. Because of restrictions on the U.S. Forest Service requiring that precise FIA coordinates be kept confidential, we downloaded image time series of Landsat and HLS and extracted point time series for each FIA site offline. Then, we calculated the two-band vegetation index (EVI2) ^57^ from the surface reflectance of Landsat and HLS time series to represent leaf greenness dynamics.

The EVI2 time series were then used to retrieve annual phenology for each FIA site using a novel Bayesian land surface phenology (BLSP) model ^58^ via the “blsp” R package ^59^. The BLSP model enables production of long-term gap-free annual phenology directly from temporally sparse Landsat observations by utilizing Bayesian methods to synthesize observations across years as prior information to help estimate phenology in each year. In this way, any specific year with limited observations benefits from borrowing information from other years for phenology retrieval. The BLSP model also quantifies uncertainty for each retrieved phenometric using a 95% credible interval (CI). To represent start-of-season (SOS) and end-of-season (EOS), we used the ‘MidGreenup’ and the ‘MidGreendown’ metrics, which correspond to the timing of inflation points of double-logistic functions fit to EVI2 time series at each pixel. The SOS and EOS inflection points correspond to the day of year when the EVI2 time series reaches 50% of its annual amplitude in spring and autumn, respectively. We chose these metrics because they are robust to noise, consistent across spatial scales, and have been widely validated using in-situ phenology records ^58,60–62^. We calculated GSL as the number of days between SOS and EOS. In total, we obtained 108,264 site-years of phenology data for 10 major species during 1985-2023 from BLSP.

To validate the BLSP retrievals, we obtained the published 30-m MSLSP30NA phenology data product (MSLSP) ^32^ from its available time span, i.e., 2016-2019 (**Extended Data Figure 2**). The ‘50PCGI’ and ‘50PCGD’ metrics in MSLSP product, representing the timing of 50% green-up and 50% green-down of the EVI2 trajectory in spring and autumn, were used as SOS and EOS, respectively. Across all site years, the coefficient of determination (R^2^) between BLSP and MSLSP were 0.87 and 0.80 for spring and autumn phenology, respectively, and the root mean square error (RMSE) values were 3.68 and 5.07 days, respectively. Within-site RMSE values were 3.0 ± 2.3 days (mean ± s.d.) for spring phenology and 4.3 ± 3.2 days for fall phenology. These results suggest that BLSP and MSLSP algorithms are highly consistent and provide confidence that phenometrics derived from the BLSP algorithm are robust and accurate.

### In-situ long-term phenology records

We used in-situ leaf phenology measurements for individual trees observed at the Harvard Forest (HF), Massachusetts, during 1990-2023 ^63^, and at the Hubbard Brook Experimental Forest (HB), New Hampshire, during 1989-2023 ^64^ (**Extended Data Figure 1**). The HF and HB phenology datasets are publicly available through their websites for each site (See Data Availability). These phenology records were tracked by in-person weekly surveys. We used the recorded timing of leaf-out in spring and leaf-fall in autumn to represent SOS and EOS, respectively; GSL was estimated as the number of days between SOS and EOS. For phenology data at HF, the timing of leaf-out is defined as the day of year on which 50% of the leaves were developed to 75% of their final size; and the timing of leaf-fall is defined as the day of year on which 50% of the leaves have fallen. For phenology data at HB, we linearly interpolated the phenology codes that describe leaf development stages in spring and autumn respectively to handle missing data, and extracted leaf-out as the day of year when a phenology code of 3 was reached in spring (leaves reach 1/2 of final length, leaves obscure half of sky as seen through crowns) and leaf-fall as the day of year on which a phenology code of 1 was reached in autumn (no more green in canopy, half of leaves have fallen, leaves still obscure half of sky as seen through crown). These phenometrics have been shown to be consistent with remotely sensed phenology ^58,60,61^. For both HF and HB phenology, trees with less than 10 years of observations were removed from the analysis.

### Climate data and trends

We downloaded the gridded Daymet meteorology dataset ^65^, which has a 1-km spatial resolution and covers North America from 1980 to present. The Daymet dataset was created by interpolating and extrapolating ground-based meteorological observations. We extracted daily temperature, precipitation, and solar radiation for each FIA site during the study period. We averaged annual and seasonal meteorology data in spring (March-May), summer (June-August), and autumn (September-November) for each year and estimated linear trends for variable during the past four decades (**Extended Data Figure 10**). These seasonal mean meteorological values were also used to quantify species-specific phenological responses to environmental factors using Bayesian hierarchical models.

### Long-term phenological dynamics

We used linear regression and the Theil-Sen slope method ^66^ to analyze long-term phenological trends. The Theil-Sen slope method was combined with the Mann-Kendall test to determine significance; these two methods are robust to outliers in trend analysis (**Extended Data Figure 3; Extended Data Figure 6**). For linear regression, we considered both the Gaussian error and the robust stand error, which accounts for heteroskedasticity (**Extended Data Figure 3**). We analyzed long-term trends of SOS, EOS, and GSL, respectively, from BLSP-derived phenometrics and from the in-situ tree-level phenology records at HF and HB.

To decompose the time series of phenometrics into interannual variation (IAV) and trend, we fit a linear regression model:

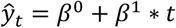

where 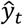 is the estimated phenology of *y*_*t*_ in year *t* for any given site; *β*^0^ is the intercept; and *β*^1^ is the trend. Then, the IAV of the phenometric is the variance of the detrended phenology:

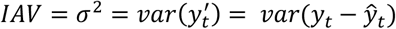

where *σ*^2^ represents variance and 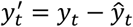 is the detrended time series. Because GSL is the duration between SOS and EOS, we have:

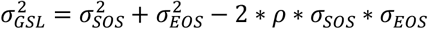

where *ρ* is the correlation between SOS and EOS. Then, to quantify the relative contributions of SOS and EOS to the IAV and trends of GSL 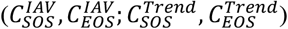, we used the following equations:

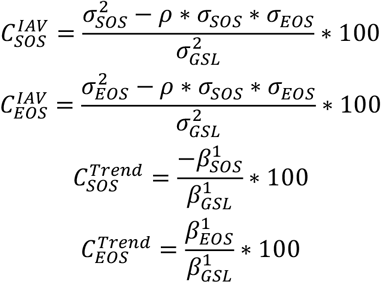

This way, for a given site, 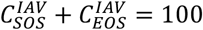 and 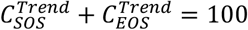 (note that the latter is an approximation because the trends were estimated rather than known).

The detrended time series 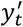 was also used to quantify the stationarity of IAV using a moving window approach. Specifically, sub-period variance (denoted by *τ*^2^) was calculated for each 9-year window across the time series, i.e., 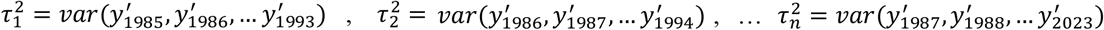. Then, a linear trend was estimated from the time series of 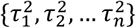 to capture the stability of the sub-period variance. If no significant trends (*p* < 0.05) were detected, the sub-period variance of the phenometric is considered temporally stable; otherwise, it experienced directional change. We tested window sizes spanning from 7 to 15 years and found the results to be similar across these different windows.

### Modeling site-specific phenology z-scores by Bayesian hierarchical models

To investigate the effects of various drivers on spring and autumn phenology across different scales, we fitted Bayesian hierarchical models to site-specific phenology z-scores, which were calculated by subtracting each site’s mean and dividing by its standard deviation. Define *n* as the total number of phenology observations (SOS or EOS) and *Y*_*ijk*_ as the observation of species *k = {1, 2, …, K}* at site *j = {1, 2, …, J}* in year *i = {1985, 1986, …, 2023}*. Then the observational model can be written as:

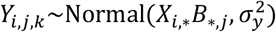

Where 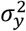 is the residual variance of the observational model; *B* is a *P* × *J* coefficient matrix where *P* is the number of predictors:

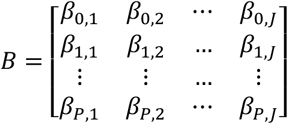

*X* is the *n* × *P* predictor matrix with the first column being ones:

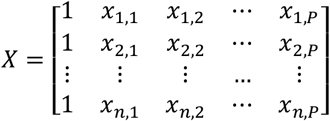

To explore the site-level effects, we added a second site-level layer to the model:

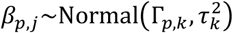

Where 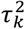 is a *K*-dimensional variance vector; Γ is a *P* × *K* matrix that stores site-level mean coefficients:

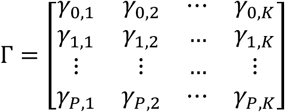

To summarize the overall effects across all species, we added a third species-level layer to the model:

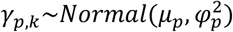

Where 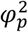 is the *p-*th element in a *P*-dimensional variance vector; *μ*_*p*_ is the *p-*th element in a P-dimensional coefficient vector. All prior distributions are non-informative, e.g., *μ*_*p*_∼Normal(0, 10000), 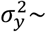*InvGamma*(0.1, 0.1).

We tested multiple meteorological variables as predictors for SOS and EOS and compared multiple model structures using site characteristics to model the sensitivity of phenology to individual meteorological variables. Model comparisons were performed using the Deviance Information Criterion (DIC) and the Watanabe-Akaike Information Criterion (WAIC). We retained the best model fit for SOS and EOS, respectively. The Bayesian model fitting was implemented using the Just Another Gibbs Sampler (JAGS, v4.3.0) and the ‘rjags’ R package.

We also tested multiple process-based phenology models but did not include the results because they involve various details in the underlying processes on which no consensus has been made and their coefficients may not provide robust site- and species-level quantifications across space ^67–72^. Also, the objective of our modeling analysis was not to find the best mechanistic model but to reveal important species-specific phenological sensitivities to environmental factors while accounting for site- and species-level variation, which can be achieved by Bayesian hierarchical modeling.

## Code and Data Availability

All R scripts used in this study will be open access via GitHub upon acceptance of the manuscript.

The extracted phenology data with true coordinates of the Forest Inventory and Analysis sites is available upon request (administrative check by U.S. Forest Service may be required due to their restrictions). The Bayesian land surface phenology (BLSP) model is open source as the ‘blsp’ R package (https://github.com/ncsuSEAL/Bayesian_LSP). Landsat and HLS observations are also open to the public through https://www.usgs.gov/landsat-missions/landsat-us-analysis-ready-data and https://hls.gsfc.nasa.gov. The long-term tree-level phenology dataset at the Harvard Forest and at the Hubbard Brook Experimental forest can be obtained from https://harvardforest1.fas.harvard.edu/exist/apps/datasets/showData.html?id=hf003 and https://portal.edirepository.org/nis/mapbrowse?packageid=knb-lter-hbr.51.14, respectively. The Daymet meteorology data can be downloaded from https://daymet.ornl.gov/.

## Acknowledgements

X.G., J.R.T., and A.D.R. acknowledge funding from the Harvard Forest Long-Term Ecological Research (LTER) grant (NSF DEB-1832210). A.D.R. also acknowledges funding from the Hubbard Brook Experimental Forest LTER grant (NSF DEB-2224545). M.A.F. acknowledges funding from the National Aeronautics and Space Administration (NASA; #80NSSC21K1974).

## Author Contributions

X.G., M.A.F., A.D.R., and J.R.T. conceptualized the research. X.G. designed the methodology and performed the analysis with resources provided by V.J.P., J.O., and A.S.B., and with feedback from M.A.F., A.D.R., and J.R.T. X.G. prepared the original draft with help from M.A.F., and M.A.F., A.D.R., J.R.T., and V.J.P. contributed to the revision.

