## Supplementary Information for "Asymmetric Spring and Autumn Phenology Control Growing Season Length in Temperate Deciduous Forests"

### Asymmetric Dynamics in Autumn Versus Spring Phenology Controls Growing Season Length in Temperate Deciduous Forests

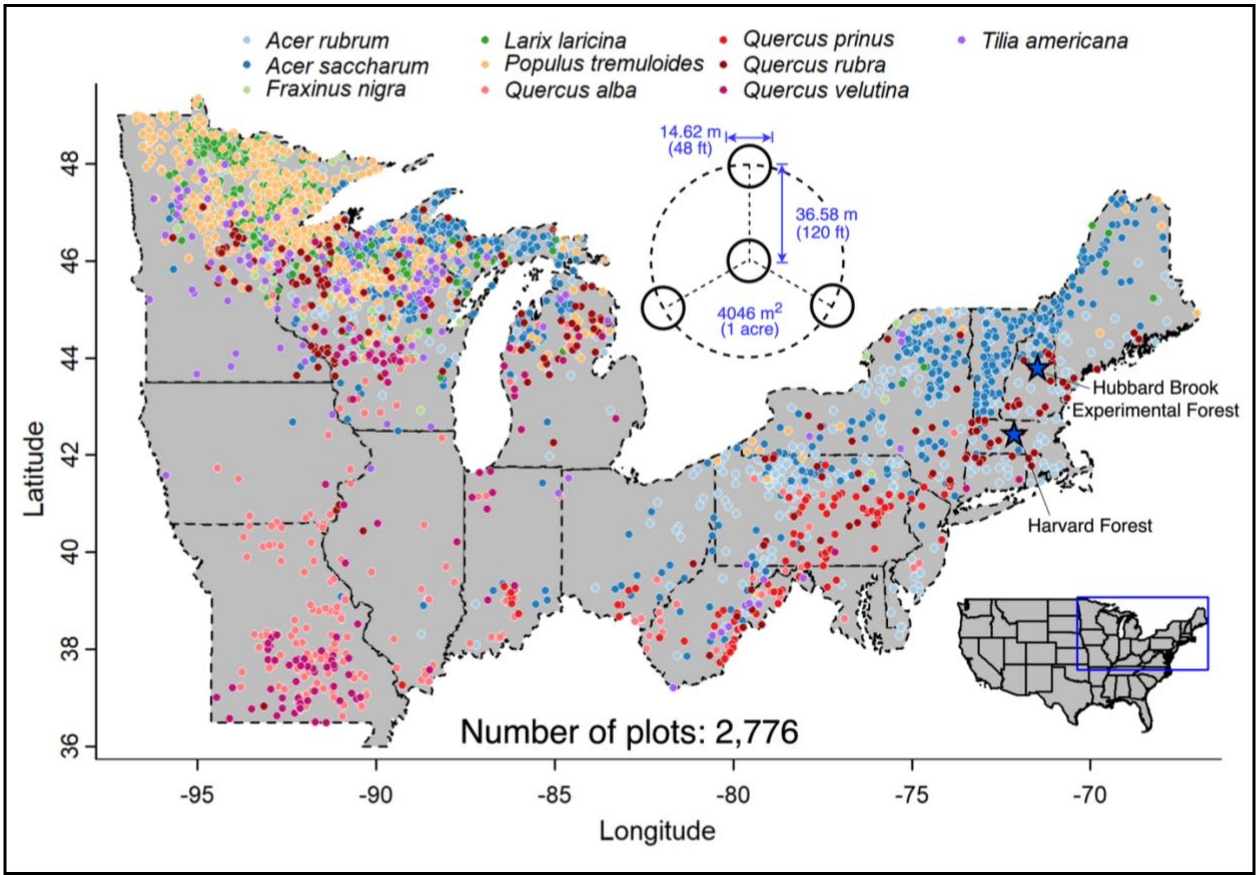


**Extended Data Figure 1 | The forest inventory and analysis (FIA) plots with dominant species in the Northeastern and Midwestern U.S.** The precise coordinates of the FIA plot locations were used in the analysis but were fuzzed here for visualization. The center inset shows the design of the FIA plot layout. The blue stars indicate the locations of the long-term in-situ phenology observed at the Harvard Forest, Massachusetts, and the Hubbard Brook Experimental Forest, New Hampshire. The corresponding species' common names and numbers of sites can be found in **Extended Data Table 1.**

**Extended Data Table 1 | Deciduous species code and common names in this study.**

| Index | Species code | Genus | Species | Leaf image | Common name | Number of sites |
|---|---|---|---|---|---|---|
| 1 | ACRU | *Acer* | *rubrum* | 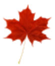 | Red maple | 374 |
| 2 | ACSA | *Acer* | *saccharum* | 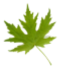 | Sugar maple | 588 |
| 3 | FRNI | *Fraxinus* | *nigra* | 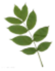 | Black ash | 118 |
| 4 | LALA | *Larix* | *laricina* | 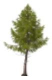 | Tamarack | 291 |
| 5 | POTR | *Populus* | *tremuloides* | 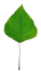 | Quaking aspen | 506 |
| 6 | QUAL | *Quercus* | *alba* | 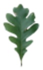 | White oak | 186 |
| 7 | QUPR | *Quercus* | *prinus* | 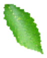 | Chestnut oak | 64 |
| 8 | QURU | *Quercus* | *rubra* | 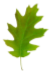 | Northern red oak | 190 |
| 9 | QUVE | *Quercus* | *velutina* | 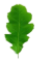 | Black oak | 99 |
| 10 | TIAM | *Tilia* | *americana* | 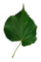 | American basswood | 103 |

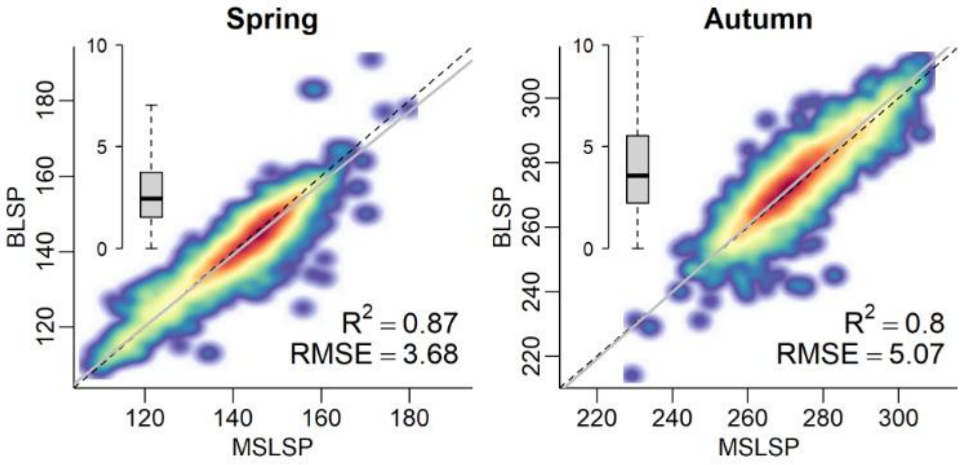


**Extended Data Figure 2 | Consistency between phenometrics of BLSP and MSLSP in MSLSP's 4-year range of 2016-2019.** The scatter plots show the comparison across all site years using a linear regression. The inset boxplots show the site-specific root mean square error (RMSE) values.

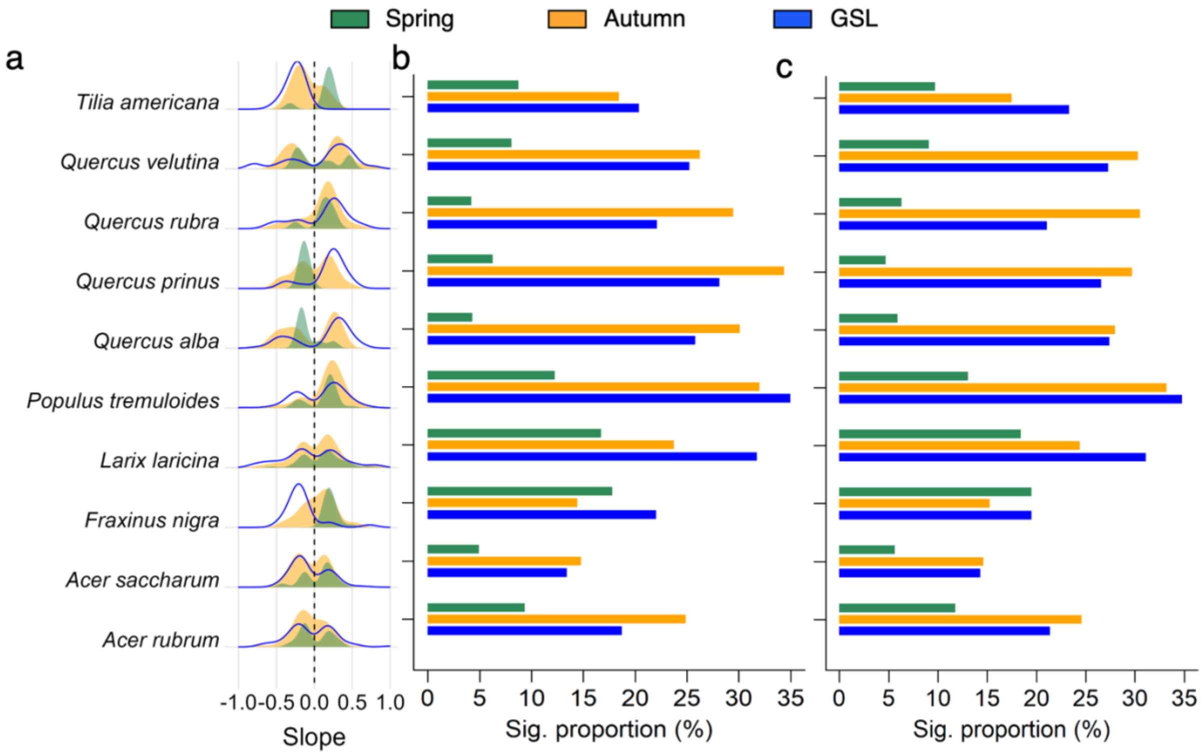


**Extended Data Figure 3 | Trend analysis same as Fig. 1 in the main text but using different approaches.** Specifically, the trends in (a) were estimated by the Theil-Sen slope and the significant proportions were derived using the Mann-Kendall test (b), which are more robust in estimating trends. Alternatively, the significant proportions were also calculated using the linear regression with robust stand errors (c), which accounts for heteroskedasticity, i.e., non-stationary variance.

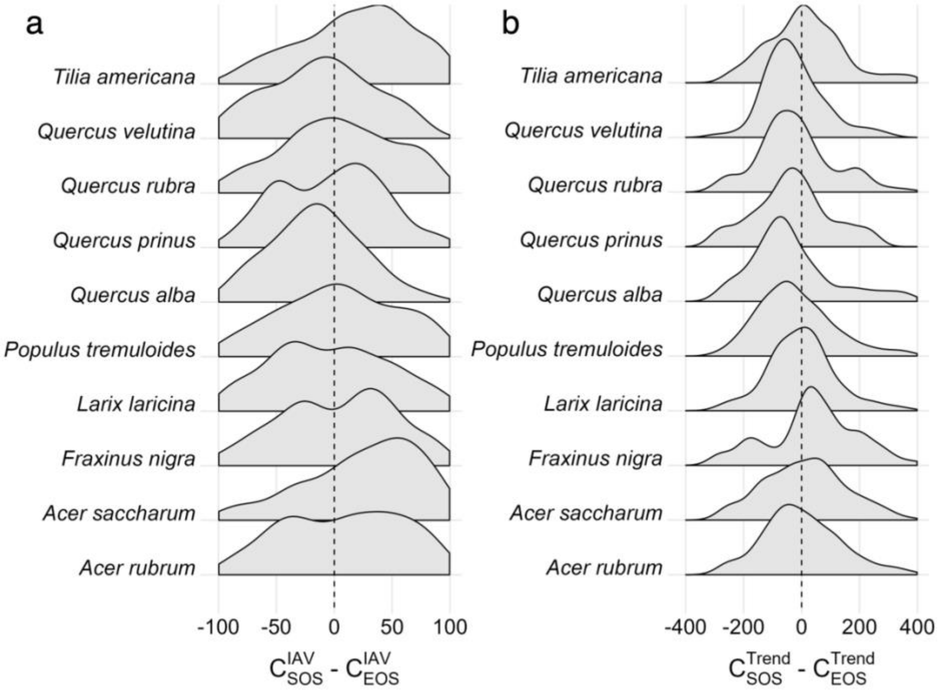


**Extended Data Figure 4 | Relative contribution of SOS and EOS to GSL in terms of interannual variation (IAV) and trend for each species and all sites.** Note that although $C_{SOS}^{Trend} + C_{EOS}^{Trend} = 100$, their values can exceed 100 or be less than 0, and thus their difference is not bounded to [-100, 100].

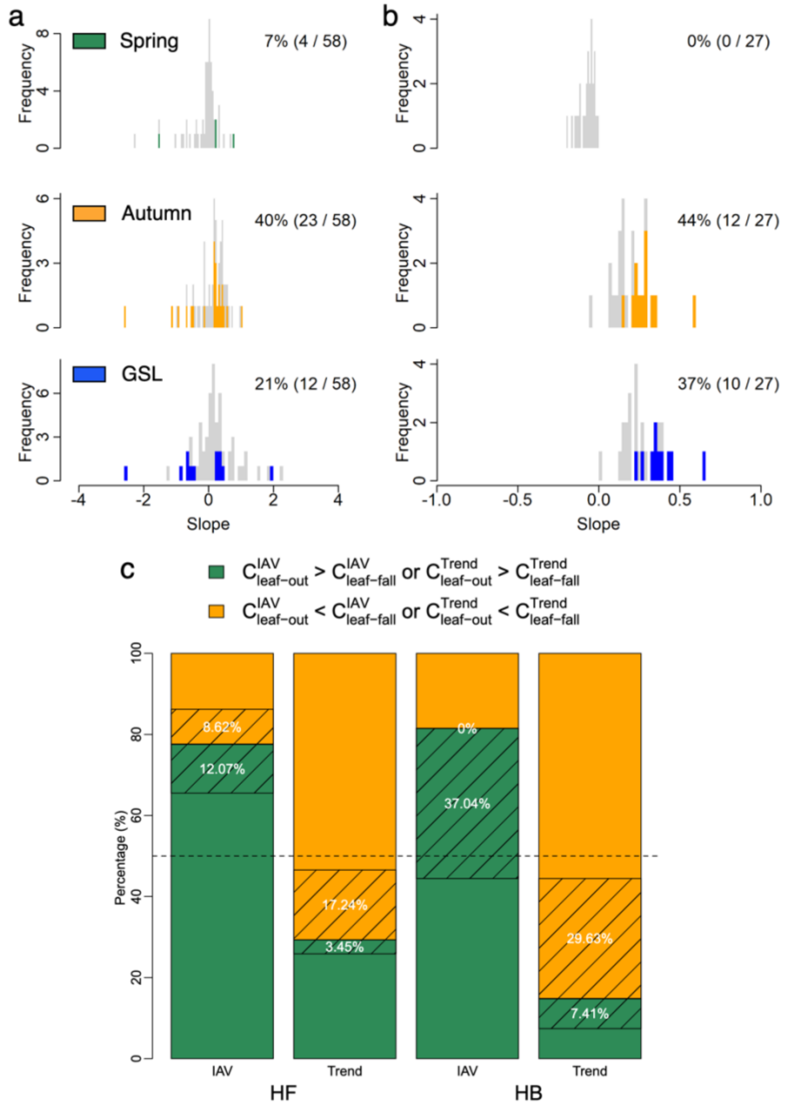


**Extended Data Figure 5 | Phenological dynamics derived from in-situ observations of individual deciduous trees.** Phenological trends observed at the Harvard Forest, Massachusetts, during 1991-2023 **(a)** and the Hubbard Brook Experimental Forest, New Hampshire, during1989-2023 **(b)**. In these datasets, spring phenology is represented by leaf-out; autumn phenology is represented by leaf-fall. Grey bars represent insignificant trends, and the colored bars represent significant trends. The labels show the significant proportions, number of significant trees, and total number of trees. **c)** The proportions of trees that had a relatively stronger leaf-out or leaf-fall contribution to GSL dynamics in terms of inter-annual variation (IAV) ($C^{IAV}_{leaf-out}, C^{IAV}_{leaf-fall}$) and trend ($C^{Trend}_{leaf-out}, C^{Trend}_{leaf-fall}$) (See Methods). The shaded rectangles and the text labels show the proportions that had a significant GSL trend. The horizonal dashed line indicates the level of 50% where leaf-out and leaf-fall had equal contributions to GSL dynamics.

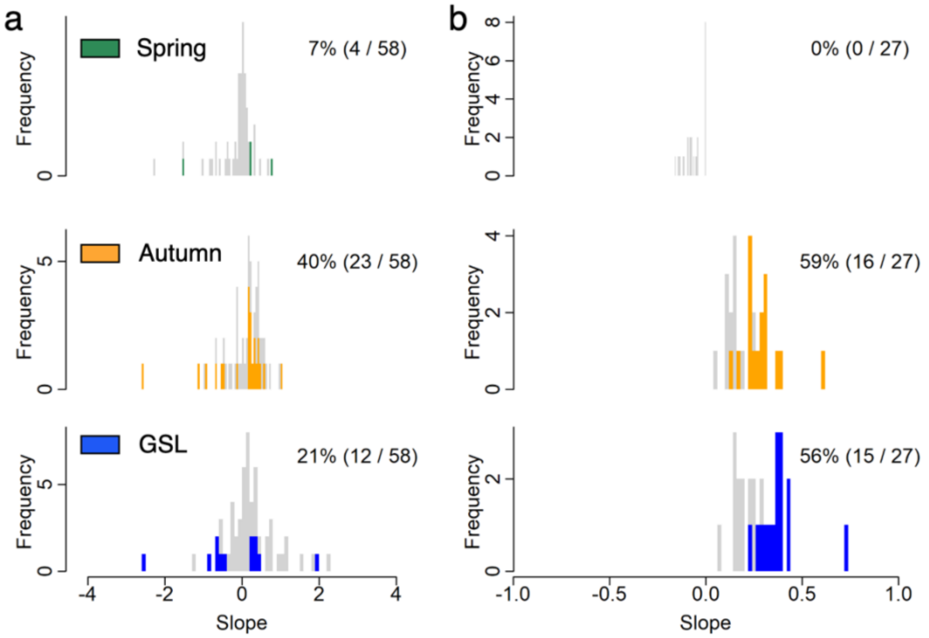


**Extended Data Figure 6 | Same as Fig. 3 but using the Mann-Kendall test as the significance test method and the Theil-Sen slope to estimate the trends.**

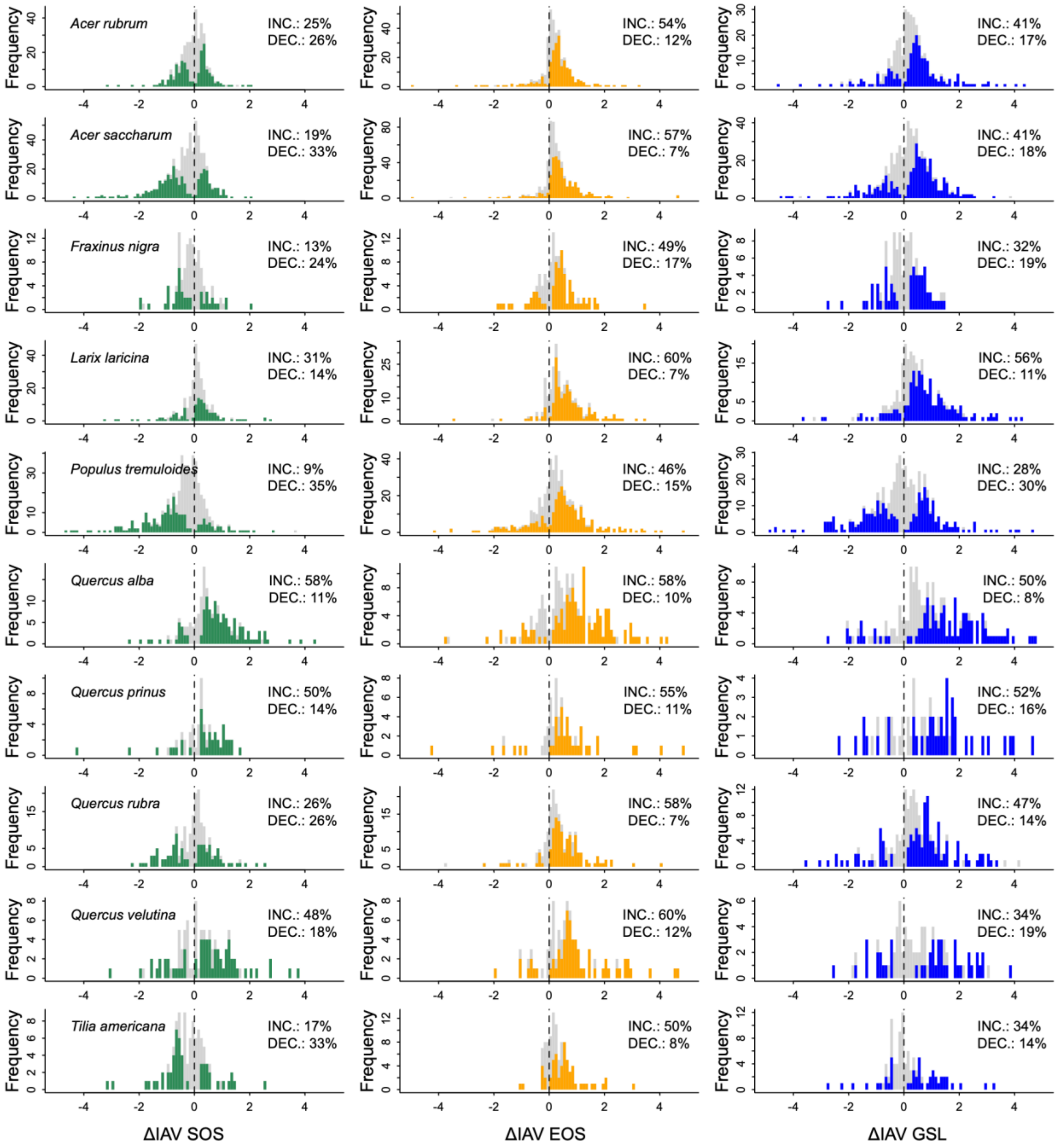


**Extended Data Figure 7 | Species-specific stationarity of phenological interannual variation (IAV).** Stationarity of IAV in SOS (ΔIAV SOS) is on the left column, EOS on the middle column, and GSL on the right column. Inner labels show the proportions of significant increasing (INC.) and decreasing (DEC.) trends. Each row represents a species, and the species name is shown on the top left side of the first column.

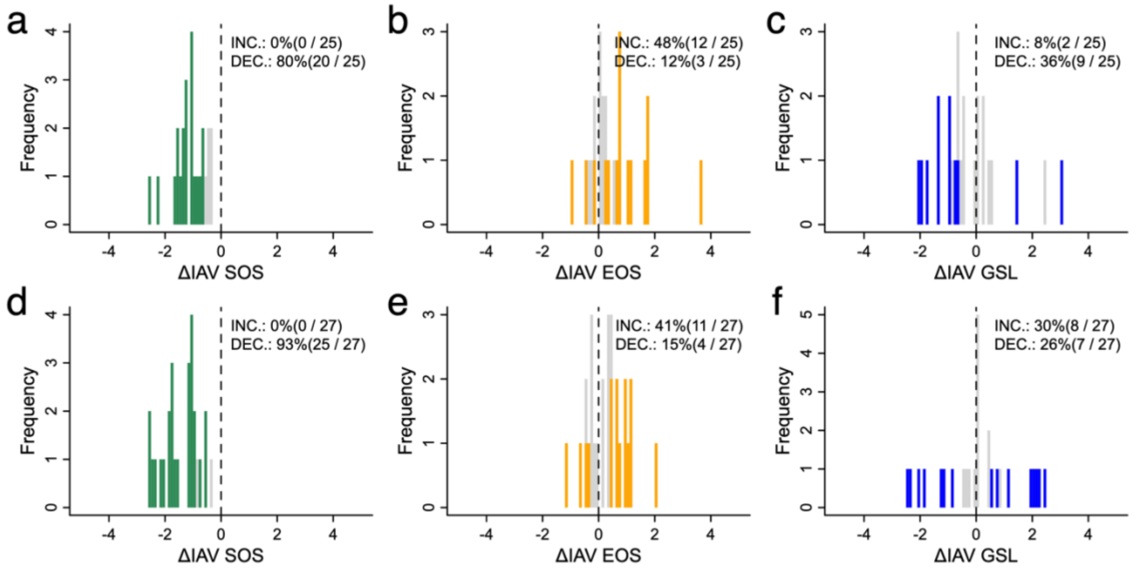


**Extended Data Figure 8 | Stationarity of phenological interannual variation (IAV) derived from ground observations.** Ground phenology observations were collected at the Harvard Forest, Massachusetts, during 1991-2023 (a-c) and the Hubbard Brook Experimental Forest, New Hampshire, during1989-2023 (d-e). Stationarity of IAV in SOS (ΔIAV SOS) is on the left column, EOS on the middle column, and GSL on the right column. Inner labels show the proportions of significant increasing (INC.) and decreasing (DEC.) trends as well as the number of trees with significant trends among the total number of trees. Trees with less than 20 years of observations were removed to accommodate the moving window approach for variance quantification.

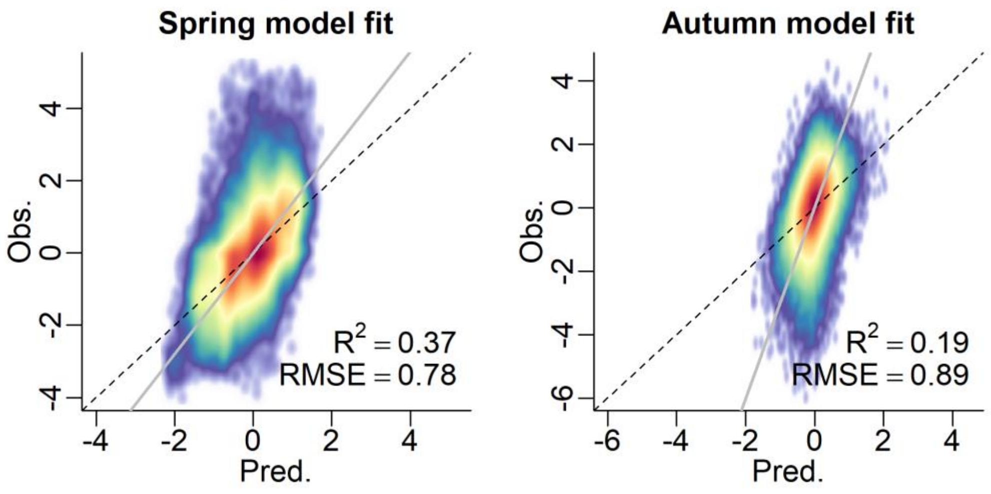


**Extended Data Figure 9 | Goodness-of-fit values of the Bayesian hierarchical models for interannual anomalies of spring and autumn phenology.**

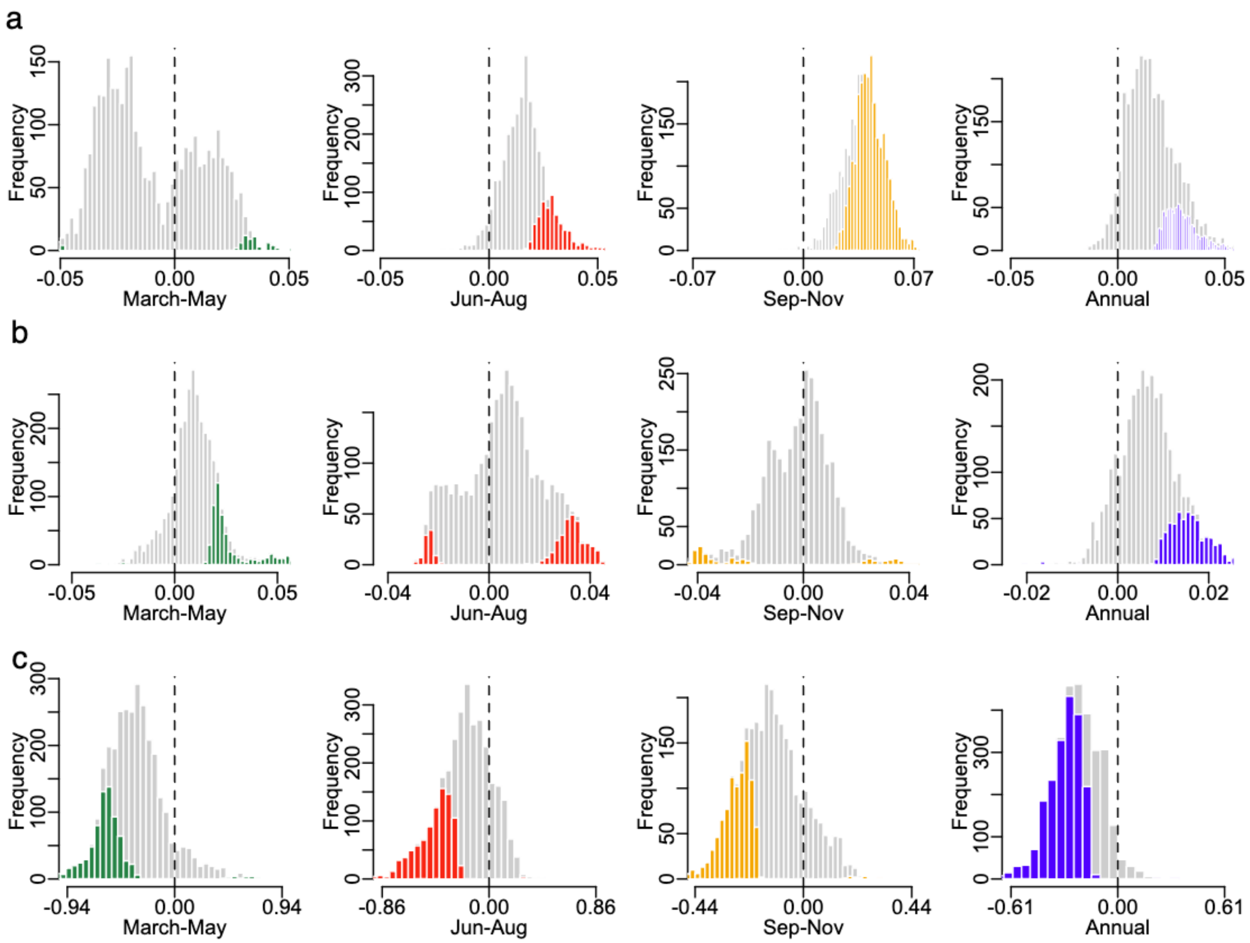


**Extended Data Figure 10 | Trends of seasonal and annual mean meteorological variables including temperature (a), precipitation (b), and solar radiation (c) for the studied sites.** The meteorological variables from 1984-2023 were obtained from the Daymet gridded climate dataset. The colored bars represent significant trends ($p < 0.05$), and the grey bars are the insignificant trends. The vertical dashed line indicates trend slope equals zero. The distinct colors represent seasons of spring (March-May; green), summer (June-August; red), autumn (September-November; yellow), and annual (blue).

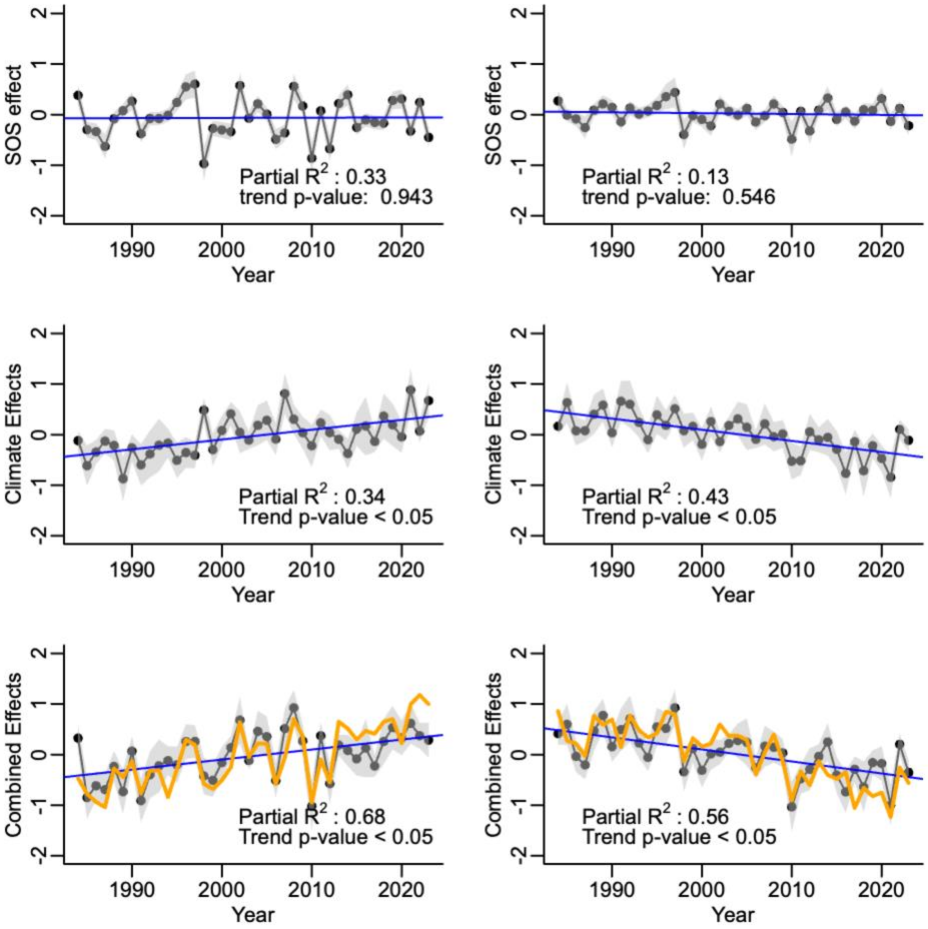


**Extended Data Figure 11 | Simulation exploring the isolated effects of SOS and climate variables on EOS trends.** The simulation was conducted by linearly regressing averaged annual z-score of EOS by predictors also in z-scores including SOS and mean temperatures, precipitation, and solar radiation in summer and autumn, respectively. The relative effects of SOS and climate variables were estimated by the model predictions with holding the other effect as the mean values. In the figure, the first column represents the dataset that was calculated from sites with no SOS trend but EOS increasing trend (319 sites). The second column from sites with no SOS trend but EOS decreasing trend (206 sites). The points are the simulated EOS values, and the blue lines represent trends. The grey polygons are confidence intervals, and the orange lines are observed mean EOS z-score values. Text labels indicate partial $R^2$ of the individual effects and p-values of the trend slopes.
